# Fish diversity drives regional productivity but not stability in southeastern United States coastal marine fishes

**DOI:** 10.1101/2021.11.10.468121

**Authors:** Juliane G Caughron, Craig J Plante, Marcel JM Reichert, Tracey I Smart, Daniel J McGlinn

## Abstract

**Aim:** Ecosystem-based management requires accurate predictions on how biotic and environmental factors interact to deliver ecosystem services. Biodiversity-ecosystem function (BEF) theory predicts that as diversity increases, the ecosystem will become more productive (positive diversity-productivity relationship – DPR) and more stable (positive diversity-stability relationship – DSR). Support for BEF has been primarily derived from fine-grained, non-harvested systems. The purpose of this study is to examine the robustness of BEF predictions for the DPR and DSR by examining how well fish diversity predicts productivity and stability of fish, shrimp, and flounder at a regional scale.

**Location:** Southeast coast of United States.

**Time Period:** 1989 - 2015

**Major Taxa Studied:** Marine Fishes

**Methods:** We used 27 years of the SEAMAP-SA Coastal Trawl Survey database to derive estimates of fish, shrimp, and flounder biomass (i.e., productivity), temporal stability of biomass (i.e., invariability of productivity), and fish community species richness. We pooled trawls into 22 km x 22 km raster cells and 3-year time bins. We controlled for variation in sampling effort using sample-based rarefaction. We compared the ability of fish species richness, water salinity, and water temperature to predict biomass and stability of all fish, shrimp, and flounder using multiple linear regression.

**Results:** Both the DPR and DSR exhibited positive log-log linear trends as expected, but the DPR had a much stronger signal. Species richness outperformed the environmental covariates in both the fish and shrimp DPR models. Surface temperature was the most important variable in both flounder models. Overall, our models better explained productivity than stability.

**Main Conclusions:** The DPR and DSR are relevant at regional scales in a commercially important fishery although support for the DSR is less justified than DPR. Further investigation into the underlying mechanisms driving the DPR and DSR are necessary to design management around BEF theory.

## Introduction

Biodiversity is declining globally (Barnosky et al., 2011), which is a major cause of concern because biodiversity influences the production and maintenance of ecosystem services that impact human health and global commercial markets (Chapin et al., 1997; Hector & Bagchi 2007; La Notte et al., 2019). Biodiversity-ecosystem function (BEF) theory suggests that if biodiversity is lost, many different ecosystem services such as biomass production and stability, pollinator services, and carbon sequestration will also be lost (Cardinale et al., 2006; Fischer et al., 2006; de Manzancourt et al., 2013).

The majority of research on BEF has focused on the diversity-productivity relationship (DPR) and the diversity-stability relationship (DSR). The DPR predicts that increasing diversity in a community will lead to increased productivity. Ecosystem productivity can be defined in various ways but most commonly it refers to total biomass in the system (Rogers et al., 2018; Tilman et al., 2014). The DSR predicts that increasing diversity in a community will increase both temporal and spatial stability (i.e., invariability; Delsol et al., 2018; Wang et al., 2017).

Although the DPR and DSR have received some general support, the majority of these studies have focused on fine-scale, terrestrial, non-harvested systems (e.g., Caldeira et al., 2005; Sanaei et al., 2018; Tilman et al., 2014; but see Gammal et al., 2019; Thibaut et al., 2012). Fine-scale studies are excellent for parsing biological mechanisms underlying BEF relationships but potentially are not relevant for regional management and policy purposes due to the scale-dependent nature of biodiversity (Bond & Chase, 2002; McGlinn & Palmer, 2009). Gamfeldt et al. (2015) carried out a meta-analysis of marine BEF experiments and found that, in general, loss of diversity results in loss of ecosystem functioning, but they noted that most of the experiments were at very fine spatial and temporal grains. Furthermore, Gonzalez et al. (2020) argued that we should expect BEF relationships to display scale dependence, which suggests that directly applying fine-scale BEF findings to the regional scale is not necessarily warranted (i.e., 10s of meters vs hundreds of km).

At the regional scale, environmental variables may play a stronger role than species richness on shaping productivity and stability (Garcia et al., 2018; Hofmann & Powell, 1998; Li et al., 2020). In the nearshore environment two key environmental parameters are temperature and salinity. Water temperature has links to energy availability and oxygen content; and is often used a geographic proxy variable (Frainer et al., 2017). Salinity is commonly used as an indicator of freshwater and estuarine inputs, and influences both resource and physiological limits of species ranges (Pecuchet et al., 2020).

Marine fish communities provide an important test case for the relevance and generality of BEF theory. Many marine fish communities have been overexploited and ineffectively managed despite their large ecological and economic importance (Rosenburg et al., 2006). This is also true of a selection of non-fish marine species that are harvested for consumption, aquaria, and production purposes (i.e., shrimps, crabs, sea slugs, oysters, scallops, etc.). Fisheries are responsible for a considerable amount of worldwide economic production and are particularly hard to study because of the difficulties of collecting data at sea at regional spatial extents. Traditionally, fisheries management has been conducted using more-mechanistic, single-species models in which each species is managed independently of their interactions with other species or their environment (Larkin, 1977; Mace, 2001). More recently, there has been a push to transition toward ecosystem-based management (EBM), with a specific directive to do so in United States fisheries in the most recent reauthorization of the Magnuson-Stevens Reauthorization Act (Magnuson-Stevens Reauthorization Act, 2007; Rosenburg et al., 2006). However, implementing EBM strategies has been slow, partly because of the significant increase in data requirements. BEF theory potentially provides an alternative basis for developing EBM strategies that harness occurrence and yield data across multiple taxa but does not require more difficult-to-collect demographic parameters required by species-specific mechanistic models.

In addition to testing the relevance of the BEF theory in a regional marine community, it is also important to understand its applicability relative to specific commercially and recreationally important groups, as it is unclear whether BEF relationships apply in contexts that are under active harvest such as marine fisheries (e.g., Frank et al., 2016). Harvest efforts are typically focused on a small suite of higher trophic-level species that may have a disproportionate effect on the total productivity of the community over other drivers such as diversity. It may be that BEF relationships such as the DPR and DSR break down in heavily exploited systems because removal of biomass by harvesters has the ability to mask a community’s intrinsic ecosystem connections. However, BEF theory has the potential to become a particularly helpful EBM tool if it can be established that biodiversity also drives ecosystem function at regional scales that are relevant to management in both harvested and non-harvested systems. Shrimp and flounder are two important fisheries in the South Atlantic region and there is a strong demand for developing methods to predict their productivity and stability over time (Atlantic States Marine Fisheries Commission, 2018; National Marine Fisheries Service, 2020). Both groups have experienced intense harvesting pressure in both commercial and recreational capacities and their respective management strategies are extremely topical in the current South Atlantic fisheries landscape (Atlantic States Marine Fisheries Commission, 2018; National Marine Fisheries Service, 2020).

The purpose of this study is to examine the efficacy of BEF predictions for the DPR and DSR using three groups of marine species (coastal fish, shrimp, and flounder) at a regional scale that is relevant for developing future policy and management. Specifically, we will examine the strength of fish species richness as a predictor of productivity and stability of productivity of all fish species, exclusively shrimp, and exclusively flounder relative to key environmental variables using a long-term fisheries-independent database in the southwestern Atlantic Ocean. We expect positive DPR and DSR relationships for both fish and flounder. It is unclear if fish species richness will serve as a good predictor of productivity and stability for another taxonomic group in the community, so we expect the DPR and DSR to be much weaker for shrimp.

## Methods

### Sampling and data acquisition

Data were acquired from the South Carolina Department of Natural Resources Southeast Area Monitoring and Assessment Program -South Atlantic (SEAMAP-SA) Coastal Trawl Survey. This ongoing survey effort began sampling in 1986 and extends from Cape Hatteras, NC to Cape Canaveral, FL. Sampling takes place on a 75 ft dual-rigged shrimp trawler during three sampling seasons (Spring, Summer, Fall) annually (see Zimney & Gut, 2020 for details).

Standardized sampling was adopted in two phases in 1989, including the establishment of the permanent strata, nets, and tow procedures in the spring and the additional establishment of daytime only sampling in the summer. The sampling area was divided into 24 strata across six regions (Zimney & Gut, 2020). Sites available to be sampled within each stratum were identified as viable trawl sites (no hangs, soft bottom). For each sampling season a random subset of the viable trawl sites within each stratum were sampled. As a result, sampling at a particular site was not continuous through time, although all strata were sampled consistently. Some sites were sampled every season while others less frequently. Sampling effort was also skewed by weather events. Hurricanes and rough seas precluded sites and even entire strata from being sampled; in particular, a single stratum in Fall 1990 and two strata in Spring 2013. A CTD was deployed at each trawl location to collect environmental data. At each site, two double-rigged trawls were conducted simultaneously and are collectively referred to as a trawl event. Specimens in the catch were then identified to the lowest taxonomic level possible, counted, and a total weight by species was determined.

### Data filtering

We filtered the data into three groups: all fish species, shrimp (*Farfantepenaeus aztecus*, *Farfantepenaeus durorarum*, *Litopenaeus setiferus*), and flounder (*Paralichthys albigutta*, *Paralichthys dentatus*, *Paralichthys lethostigma*). All fish were recorded at the species level except for jawfishes (Family Opistognathidae), scup (*Stenotomus* spp.), and mojarritas (*Eucinostomus* spp.). Anchovies (*Anchoa* spp.) were recorded at both the species and genus level and so were grouped as the genus *Anchoa*. The subset of flounder species were also included in the all fish species group; however, shrimp species were not. Fish were chosen because they comprise the majority of the total catch biomass. Shrimp also are large contributors to catch biomass as well as being commercially important species (National Marine Fisheries Service, 2020). Flounder make up a smaller portion of catch biomass but are of interest for both harvest and management (National Marine Fisheries Service, 2020). These groups were subset to the collection years 1989 – 2015. Sampling during this timeframe was almost entirely consistent in effort and methods and allowed for even three-year time bins. One exception was spring 1989, which utilized night-time sampling while all other years and seasons utilized daytime-only sampling. Prior to 2000, samples also were taken in deep-water strata (> 30 ft). Samples from these outer strata were omitted due to differences in primary habitat type and community composition, as well as incomplete sampling over the extent of the timeframe.

### Sampling Grid

We binned the trawl events into 70 square 22.2 km x 22.2 km raster regions in three-year blocks. The spatial and temporal grain were chosen to achieve good spatial and temporal coverage while maintaining reasonable statistical power (Spake et al., 2021). Additionally, coarser temporal grains help to account for species with low detectability compared to using individual stations or strata. For a raster region to be included in the analysis it had to have a minimum of five trawl events in all nine of the three-year time bins. This minimum threshold resulted in 36 raster regions covering an estimated area of 17,424 km^2^ across 27 years. These raster regions were spread throughout the extent of the sampling area (Figure 1).

**Figure 1 -.**
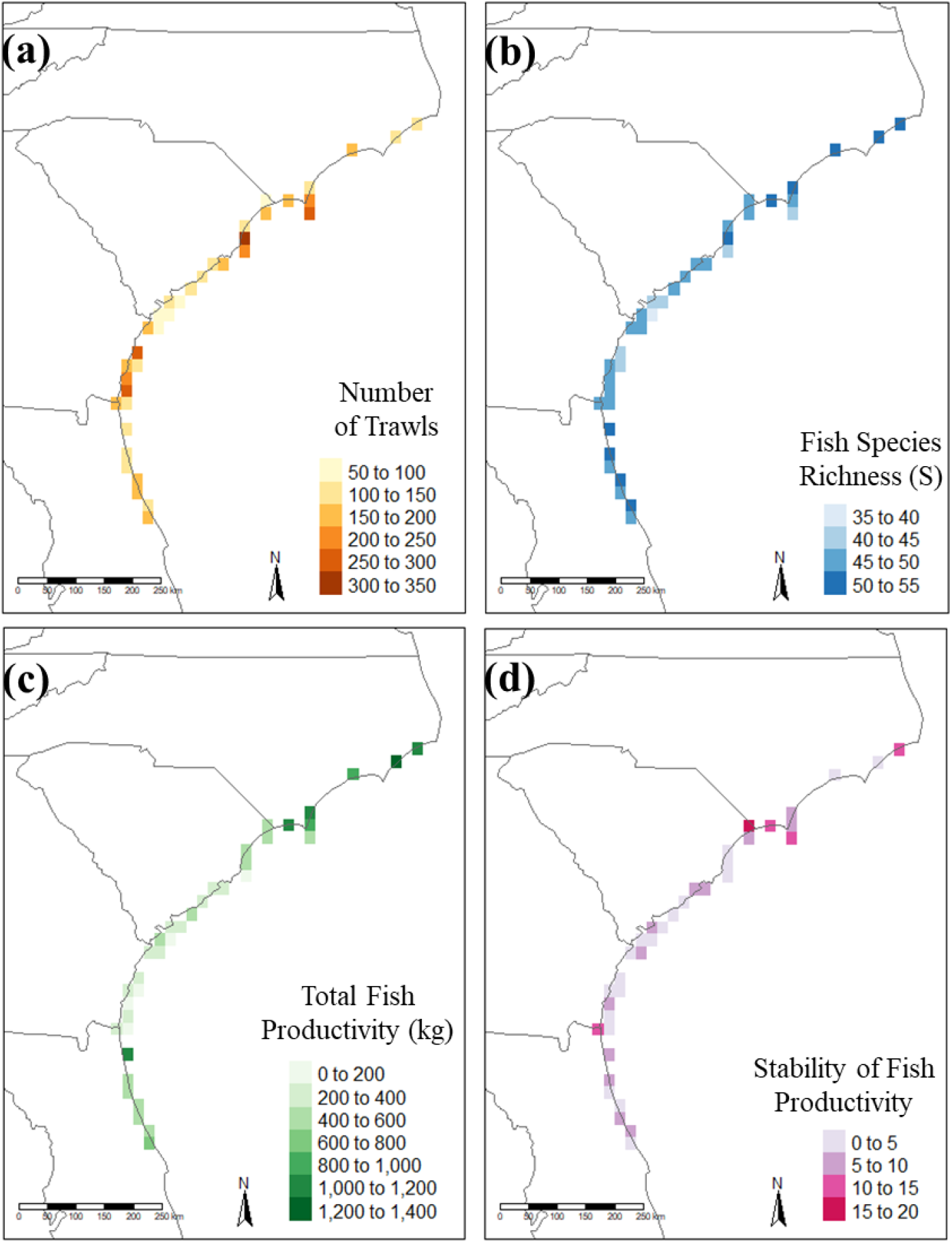
Map of 36 raster regions meeting sampling requirements spread throughout the survey extent. Raster squares are approximately 22.2 km by 22.2 km. Panels b, c, and d were rarefied to 5 trawl event subsamples due to unequal effort. a) Total number of trawls in each raster region through time illustrates unequal effort in certain areas. b) Average fish species richness per raster region across nine time bins. c) Average fish total biomass (kg) per raster region across nine time bins. d) Average temporal stability of fish biomass per raster region across nine time bins.

### Calculating species richness, biomass, and stability of biomass post-rasterization

To correct for unequal trawl event density across the raster regions (Figure 1a), we randomly subsampled each raster, time-bin to 5 trawl events (i.e., sample-based rarefaction). Each metric we examined was computed as the average of 1,000 bootstrap iterations that each drew a different random subset of trawl events. Cumulative species richness and total biomass were calculated within each raster and each time bin. We used wet weight (kg) biomass for five random trawls as a proxy of productivity for fish, shrimp, and flounder. A community was considered to have high stability in productivity when the temporal variance in total biomass was low. Temporal stability of biomass was calculated using the invariability (I(A) = mean^2^(productivity of an area) / variance(productivity of an area)) metric, defined in Wang et al. (2017) and Delsol et al. (2018), across the time bins per raster. This process was repeated for all three groups: fish, shrimp, and flounder.

### Biomass, stability, and species richness relationships

We used multiple linear regression models to predict productivity and stability using species richness, surface water temperature, and surface salinity as explanatory variables. As noted in the introduction, surface temperature was used as a proxy for oxygen availability and physiological processes, and surface salinity was a proxy of freshwater and estuarine inputs. Biomass, stability, and species richness were log_2_-transformed prior to model fitting and standardized beta coefficients reported. We used lowess smoother functions to examine our assumptions of linearity.

## Results

### Geographic Trends

Overall fish species richness (Figure 1b), biomass (Figure 1c), and stability (Figure 1d) appeared to display nonrandom geographic structure. There were two areas of elevated species richness, biomass, and stability near the border of North Carolina and South Carolina and off northern Florida (Figure 1b-d).

Surface water temperature was highly correlated with latitude with the lowest temperatures in the north, gradually increasing moving south (Figure 2a). Surface salinity was lowest in the middle of the sampling range off the coast of South Carolina and Georgia (Figure 2b), which is indicative of greater freshwater input in these regions from nearby estuarine systems.

**Figure 2 -.**
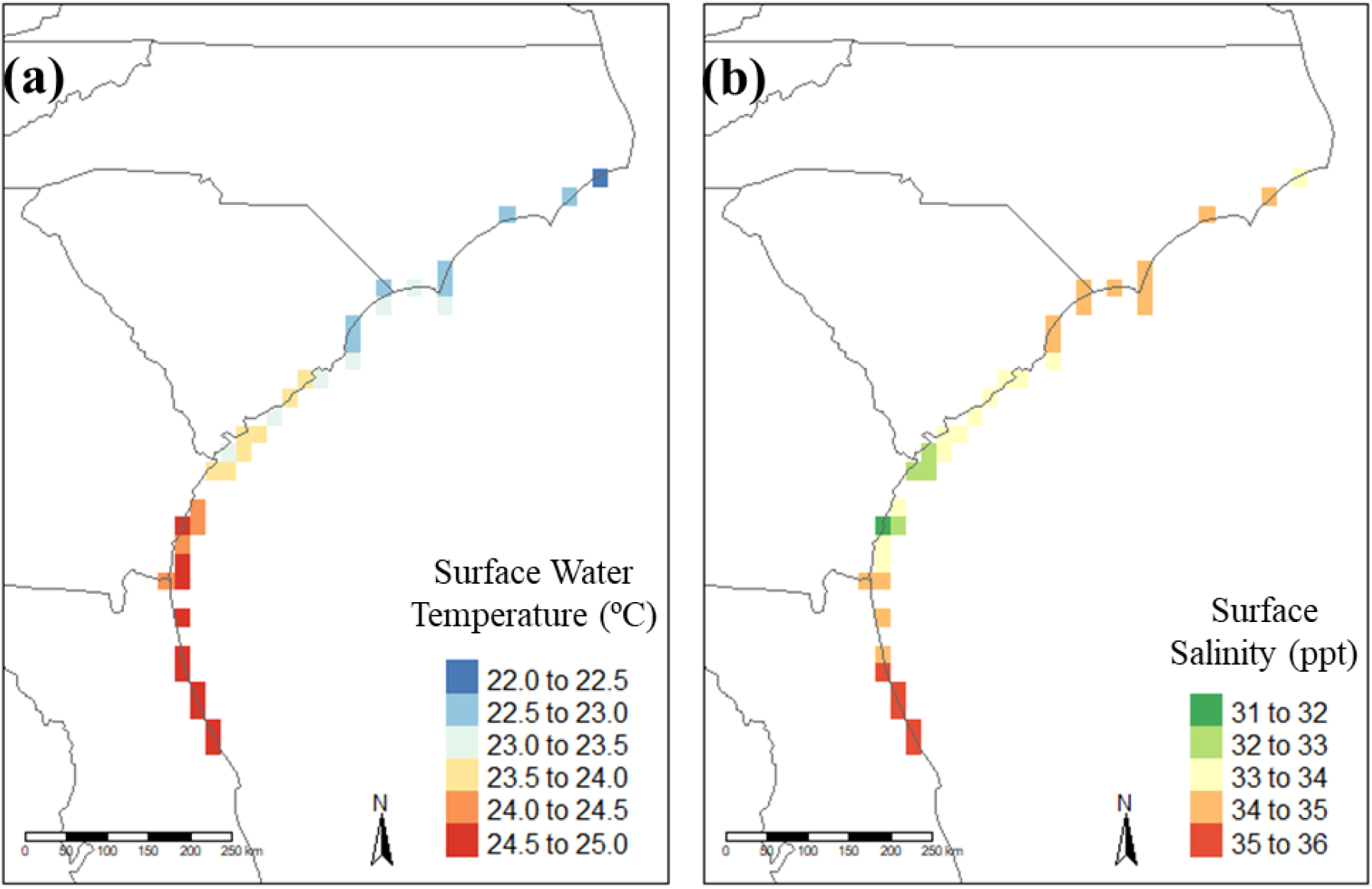
Map of average surface temperature and average surface salinity throughout the 36 raster regions. Raster squares are approximately 22.2 km by 22.2 km. a) Average surface temperature (°C) per raster region through time. b) Average surface salinity (ppt) per raster region through time.

### Productivity and Stability Models

#### Fish

Our models captured more of the variation in biomass (adjusted *R*^2^ = 0.79) than in stability (adjusted *R*^2^ = 0.07). Productivity and stability were both positively correlated with species-richness (Figure 3, Table 1). Additionally, we found that species richness was more important than temperature or salinity for predicting productivity and stability (Figure 3, Table 1). Surface temperature was negatively related to productivity but positively related to stability (Figure 3, Table 1). Surface salinity was a better predictor of stability than of productivity (Figure 3, Table 1). Residuals of neither the DPR or DSR models appeared to be spatially structured (Supplemental Figure S1).

**Figure 3 -.**
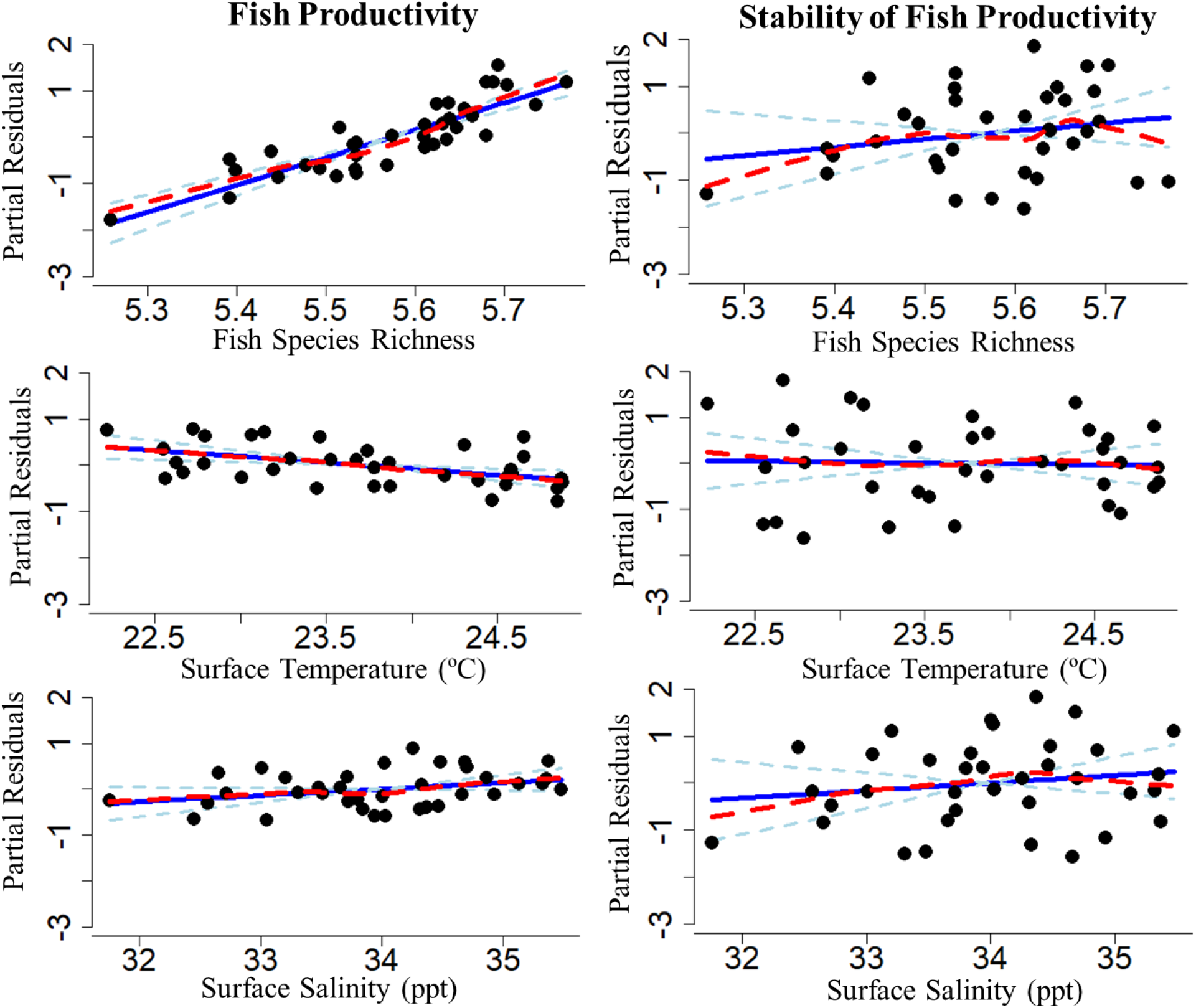
Partial residuals of log_2_(fish productivity) and log_2_(stability of fish productivity) relative to log_2_(fish species richness), surface temperature, and surface salinity after controlling for the other explanatory variables. The linear models are represented by the solid blue lines. The dashed light blue lines represent standard error estimates. The red dashed lines are lowess smoothers.

**Table 1 -.** Partial standardized beta coefficients (SE) and variance explained for the fitted models (Supplemental Table S1). The degrees of freedom for all models are 32. Bolded values indicate significance (p < 0.05).

| Group | Modeled Relationship | Response Variable | Standardized beta coefficients (+/- SE) for Explanatory Variables |  |  | Variance Explained |
| --- | --- | --- | --- | --- | --- | --- |
|  |  |  | *log Species Richness* | *Surface Temperature* | *Surface Salinity* | *Adjusted R<sup>2</sup>* |
| Fish | DPR | *log Productivity* | **0.75** (0.09) | **-0.25** (0.09) | 0.14 (0.08) | 0.79 |
|  | DSR | *log Stability* | 0.20 (0.19) | -0.03 (0.17) | 0.16 (0.19) | 0.01 |
| Shrimp | DPR | *log Productivity* | **0.60** (0.16) | **0.39** (0.15) | **-0.36** (0.16) | 0.29 |
|  | DSR | *log Stability* | 0.26 (0.19) | 0.06 (0.17) | -0.33 (0.19) | 0.02 |
| Flounder | DPR | *log Productivity* | **0.34** (0.14) | **-0.59** (0.12) | 0.03 (0.13) | 0.50 |
|  | DSR | *log Stability* | -0.13 (0.18) | **-0.41** (0.16) | -0.12 (0.18) | 0.12 |

#### Shrimp

As with all fish, our explanatory variables explained more variation in shrimp productivity (adjusted *R*^2^ = 0.29) than in shrimp stability (adjusted *R*^2^ = 0.02). Shrimp productivity was positively correlated with both fish species richness and surface water temperature (Figure 4, Table 1). There was a weakly positive relationship between shrimp stability and both fish species richness and surface water temperature (Figure 4, Table 1). Conversely, surface salinity was negatively correlated with both shrimp productivity and stability (Figure 4, Table 1). Similar to the fish group, residuals of neither the DPR or DSR models appeared to be spatially structured (Supplemental Figure S1).

**Figure 4 -.**
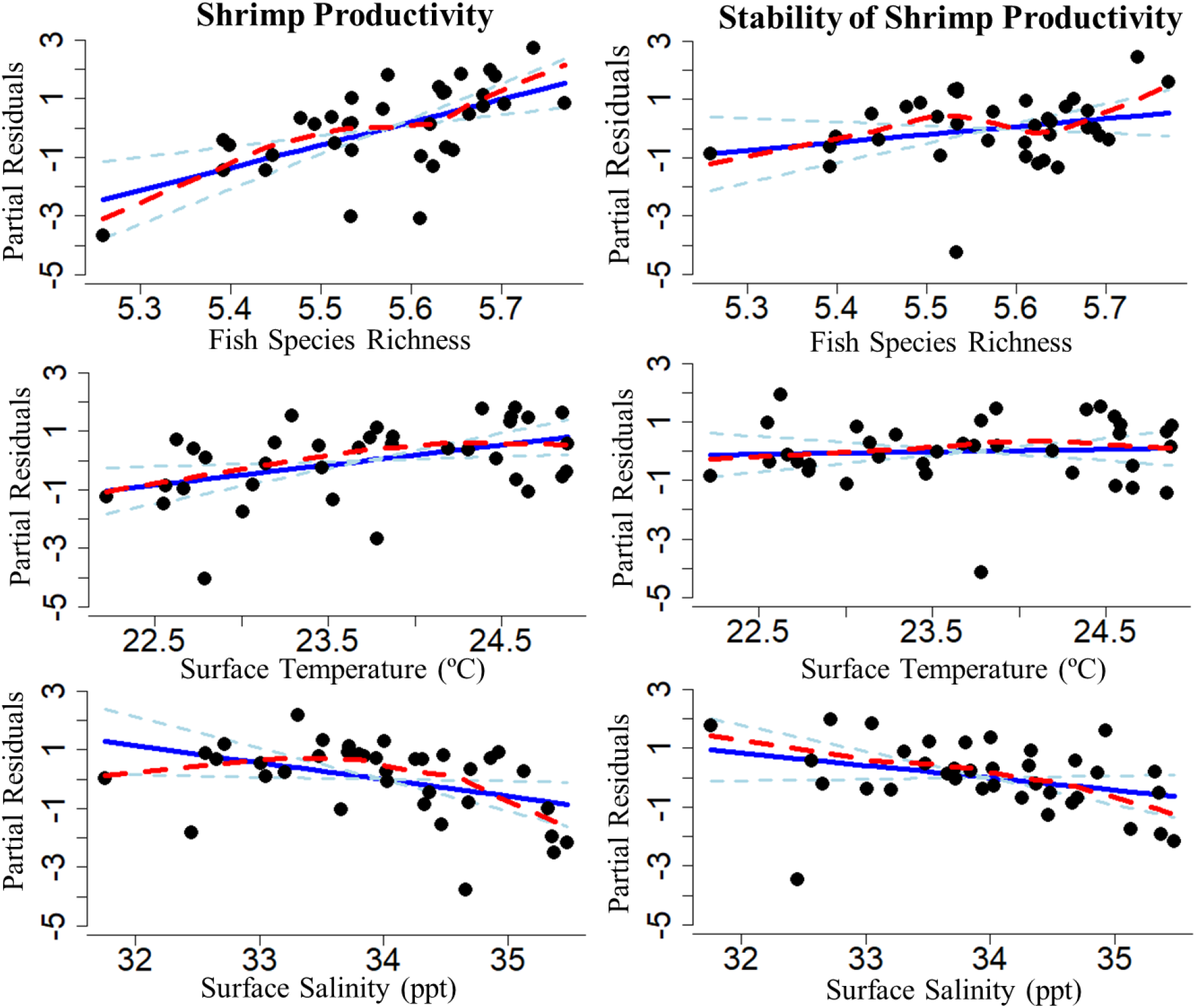
Partial residuals of log_2_(shrimp productivity) and log_2_(stability of shrimp productivity) relative to log_2_(fish species richness), surface temperature, and surface salinity after controlling for the other explanatory variables. The linear models are represented by the solid blue lines. The dashed light blue lines represent standard error estimates. The red dashed lines are lowess smoothers.

#### Flounder

Again, the flounder productivity model (adjusted *R*^2^ = 0.50) captured more variation than its stability counterpart (adjusted *R*^2^ = 0.12). Fish species richness was positively correlated with flounder productivity (Figure 5, Table 1). Fish species richness was neither positively nor negatively correlated with flounder stability. The same was true of surface salinity relative to flounder productivity or flounder stability (Figure 5, Table 1) Surface water temperature exhibited a negative correlation with both flounder productivity and stability (Figure 5, Table 1). Once more, residuals of neither the DPR or DSR models appeared to be spatially structured (Supplemental Figure S1).

**Figure 5 -.**
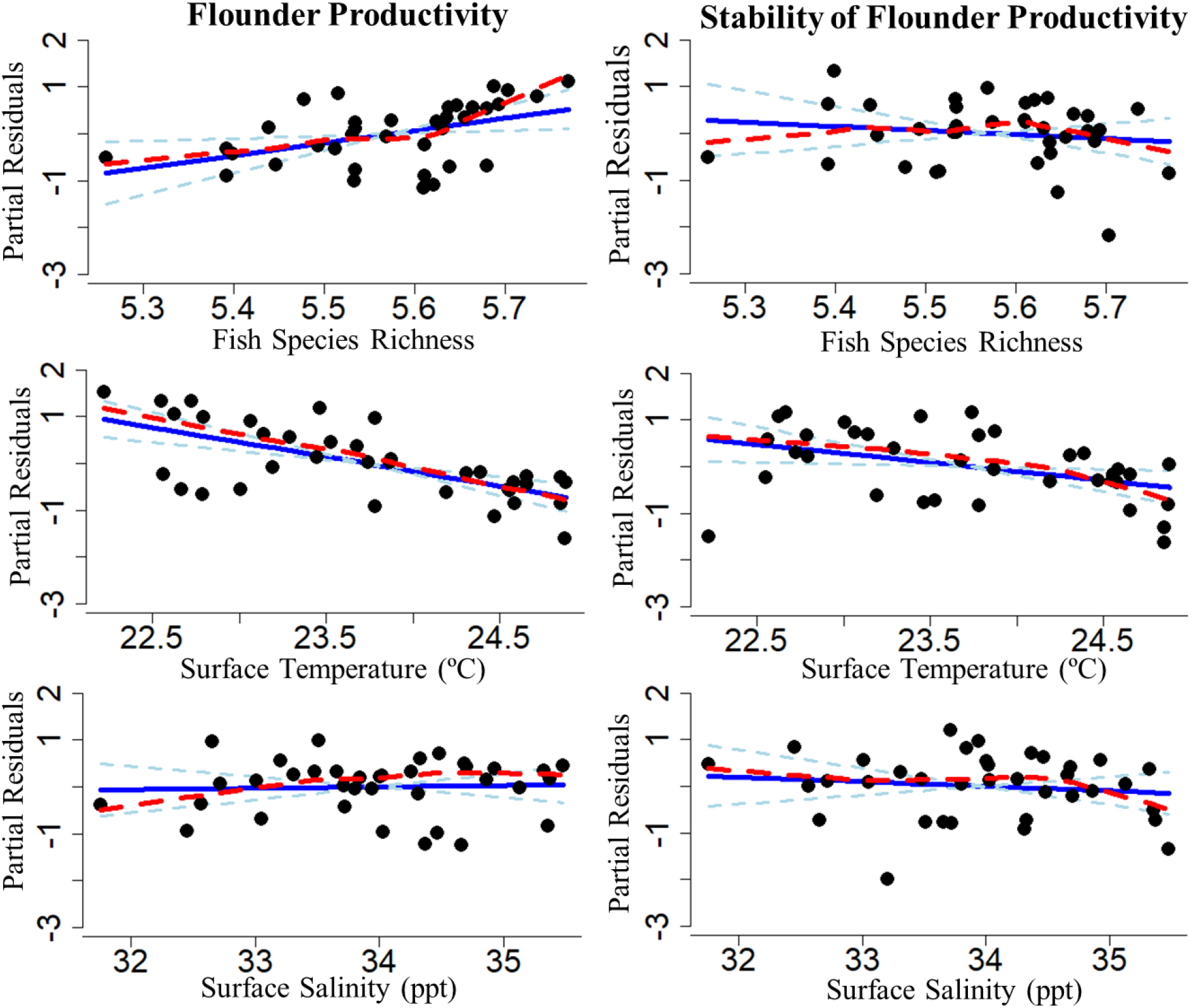
Partial residuals of log_2_(flounder productivity) and log_2_(stability of flounder productivity) relative to log_2_(fish species richness), surface temperature, and surface salinity after controlling for the other explanatory variables. The linear models are represented by the solid blue lines. The dashed light blue lines represent standard error estimates. The red dashed lines are lowess smoothers.

## Discussion

We explored two key biodiversity-ecosystem function (BEF) relationships in a regional-scale marine context. We found that areas with higher fish diversity had higher fish productivity (i.e., a positive DPR), but we did not find that diversity was linked with ecosystem stability (i.e., there was no relationship in the DSR). Additionally, fish species richness was a better predictor of productivity than the environment (temperature or salinity). Our results suggest that the DPR, but not necessarily the DSR, is relevant at regional extents in fish communities under active recreational and commercial harvesting. This is important because the majority of work on BEF relationships has focused on fine grains and small scales in terrestrial, non-harvested communities (Caldeira et al., 2005; Tilman et al., 2014). Our findings suggest that scaling up BEF relationships from fine-grained and fine-scaled experiments is not necessarily straightforward, particularly when trying to predict ecosystem stability (Bond & Chase, 2002; Gonzalez et al., 2020; Thompson et al., 2018; Zhang et al., 2018). Our results on the DPR are consistent with fine-scaled experimental (Gamfeldt et al., 2015) and broad-extent observational (Lefcheck et al., 2019; Thibaut et al., 2012) marine studies that also found positive relationships between species richness and ecosystem functioning.

We found that fish diversity had some utility in predicting the productivity but not stability of economically relevant sub-groups such as shrimp and flounder. Shrimp productivity was positively related to fish diversity, but temperature and salinity were also strong covariates. Flounder productivity was also positively related to fish diversity, but temperature had a much stronger effect. Fish diversity did not covary with the stability of either sub-groups. BEF theory traditionally has been tested in taxonomic groups that are not under intense pressure from commercial or recreational exploitation. Our results demonstrate that BEF theory can provide a first approximation of yield of economically important groups. However, we suggest that as the types of species considered in the yield calculation narrows, more traditional biologically informed models (e.g., ones that incorporate organism physiology or life history for the specific group of interest) likely will be needed to provide appropriate management guidance (e.g., Aeberhard et al., 2018).

We observed a strong DPR, but the causal directionality between diversity and productivity is still unclear. It may be that areas with higher species diversity promote productivity due to niche partitioning (e.g., Cardinale et al., 2012; Isbell et al., 2018) or/and it may be that more productive regions support higher diversity due to resource constraints (e.g., Wright, 1983; Storch & Okie, 2019). Gross & Cardinale (2007) developed models which demonstrated that both causal directions can exist simultaneously at a particular spatial scale. Gamfeldt et al. (2015) synthesized the results of 110 BEF marine experiments which purposely manipulated diversity; and found that, in general, richness was positively linearly related to increased productivity in polycultures relative to monocultures. Thus, there is support that in marine ecosystems diversity can drive productivity, but Gamfeldt et al. (2015) noted that most of their studies were at very fine spatial and temporal scales and therefore had questionable relevance at regional scales. Using an observational study, Craven et al. (2020) found equal empirical support across a large number of forest inventory plots for both directionalities of the DPR in North American forests.

It is also important to recognize that the DPR may be driven by an artefact in which another process is responsible for driving both diversity and productivity and that these variables have no direct causal link. In our study region, we observed that biomass, species richness, and stability were higher near the border between North Carolina and South Carolina (Figure 1). This hotspot is relatively near the Gulf Stream and has been described previously as a biogeographic and phylogeographic barrier (Baker et al., 2008; Calder et al., 1992; Engle et al., 1999; Rodriguez-Rey et al., 2017). It could be that the biogeographic setting rather than diversity per se is what is driving high productivity and high diversity in this area. Further research and novel methods are needed to disentangle the causality underlying the DPR at regional scales where experimental approaches are untenable. However, regardless of the exact causality, the existence of a strong DPR is a useful tool for developing regional management and conservation priorities.

An important next step towards better understanding the mechanisms underlying BEF relationships is to investigate the degree to which spatial and temporal interspecific asynchrony contributes to the DPR and DSR (Loreau & de Mazancourt, 2013). Two key mechanisms that may contribute to increased stability in an asynchronous system is the variation in the strength at which species react to an environmental driver and the differences in the rate at which species respond to perturbations (Loreau & de Mazancourt, 2013; Yan et al., 2021). Another avenue that warrants additional investigation is how the particular metrics of productivity and stability used may influence these relationships. In this study, we used total biomass as a proxy for ecosystem productivity, but biomass turnover, consideration of commercial value and other factors may also be important in evaluating ecosystem productivity (Clausen et al., 2017; Morais & Bellwood, 2020). Assessing the role these factors play in driving the shape and strength of these patterns will likely be rather beneficial in understanding DPR and DSR mechanisms. It may also be important to consider not only the total number of species within a defined area but also how species composition changes throughout the landscape, and if considering beta diversity or other diversity metrics deliver the same DPR and DSR patterns observed here (e.g., Omidipour et al., 2021).

Additionally, it is necessary to extend BEF theory so that it can provide real management applications with actionable outcomes. There are a few specific management scenarios we feel would benefit from DPR- and DSR-driven decision making. One such scenario is in the case of a data-poor community and/or stock. Species management plans and stock assessments are particularly data-intensive and are hard to enact in data-poor conditions. By encouraging management that promotes and maintains high species richness, a DSR paradigm may serve as a placeholder management plan that could be enacted in the short term to promote stability in the habitat while the necessary data for a more extensive long-term management plan is being collected.

DPR and DSR models can also help foster a transition from single-species mechanistic models to ecosystem-based management (EBM) in the fisheries sector. Monitoring abundances and the covariance matrices of a community to manage for production and stability may serve as an effective EBM tool. The DPR and DSR may also help in identifying effective MPA locations that are or were productive and stable ecosystems. These might be given priority when choosing where extra protections from fishing and other exploitation are enacted.

BEF theory has primarily been focused on small-scale terrestrial experiments with only a select number of examples from either larger scales or marine environments (Gamfeldt et al., 2008; Stachowicz et al., 2007; Thibaut et al., 2012). Expanding this sector of biodiversity research to a larger-scale, commercially relevant context such as marine fish is imperative for developing useful management applications. Further research regarding the mechanisms underlying BEF relationships, as well as the scaling of the DPR and DSR, are important in continuing to bridge the gap between understanding these ecological phenomena and providing advice and tools for managers. Ecosystem-based management efforts in fisheries will benefit from a better understanding of the relevance and underpinnings of BEF models in marine ecosystems.

## Acknowledgements

The authors would like to thank members of the SEAMAP-SA Coastal Trawl Survey for their helpful discussions and expertise relative to the survey and data set. This project was funded in part by the Joanna Foundation Graduate Fellowship in Marine Biology.

## Biosketch

Juliane Caughron is a quantitative fisheries ecologist interested in spatial, movement, and diversity patterns and how these topics can better inform interspecific and ecosystem-based management strategies.

## Data Accessibility Statement

Data from the SEAMAP-SA coastal trawl survey can be accessed at https://www2.dnr.sc.gov/seamap/. All R code for analysis can be accessed at https://github.com/mcglinnlab/fish_stability. Upon acceptance code will be submitted to zenodo.

