## Supplemental Materials for "Fish diversity drives regional productivity but not stability in southeastern United States coastal marine fishes"

**Supplemental Table S1**- Raw coefficients (+/- SE), intercept, and variance explained for the fitted models. The degrees of freedom for all models are 32. Bolded values indicate significance ($p<0.05$).

| Group | Modeled Relationship | Response Variable | Raw coefficients (+/- SE) for Explanatory Variables | | | | Varience Explained |
|---|---|---|---|---|---|---|---|
| | | | *log Species Richness* | *Surface Temperature* | *Surface Salinity* | *Intercept* | *Adjusted $R^2$* |
| **Fish** | DPR | *log Productivity* | **5.876 (0.682)** | **-0.265 (0.085)** | 0.138 (0.082) | **-22.44 (4.335)** | 0.7913 |
| | DSR | *log Stability* | 1.723 (1.610) | -0.037 (0.200) | 0.163 (0.193) | -12.158 (10.226) | 0.0121 |
| **Shrimp** | DPR | *log Productivity* | **7.776 (2.095)** | **0.690 (0.261)** | **-0.571 (0.251)** | **-37.356 (13.310)** | 0.2923 |
| | DSR | *log Stability* | 2.719 (1.999) | 0.081 (0.249) | -0.424 (0.239) | -1.952 (12.699) | 0.01561 |
| **Flounder** | DPR | *log Productivity* | **2.660 (1.051)** | **-0.627 (0.131)** | 0.0298 (0.126) | -0.614 (6.674) | 0.504 |
| | DSR | *log Stability* | -0.877 (1.234) | **-0.383 (0.153)** | -0.100 (0.148) | **18.834 (7.838)** | 0.1213 |

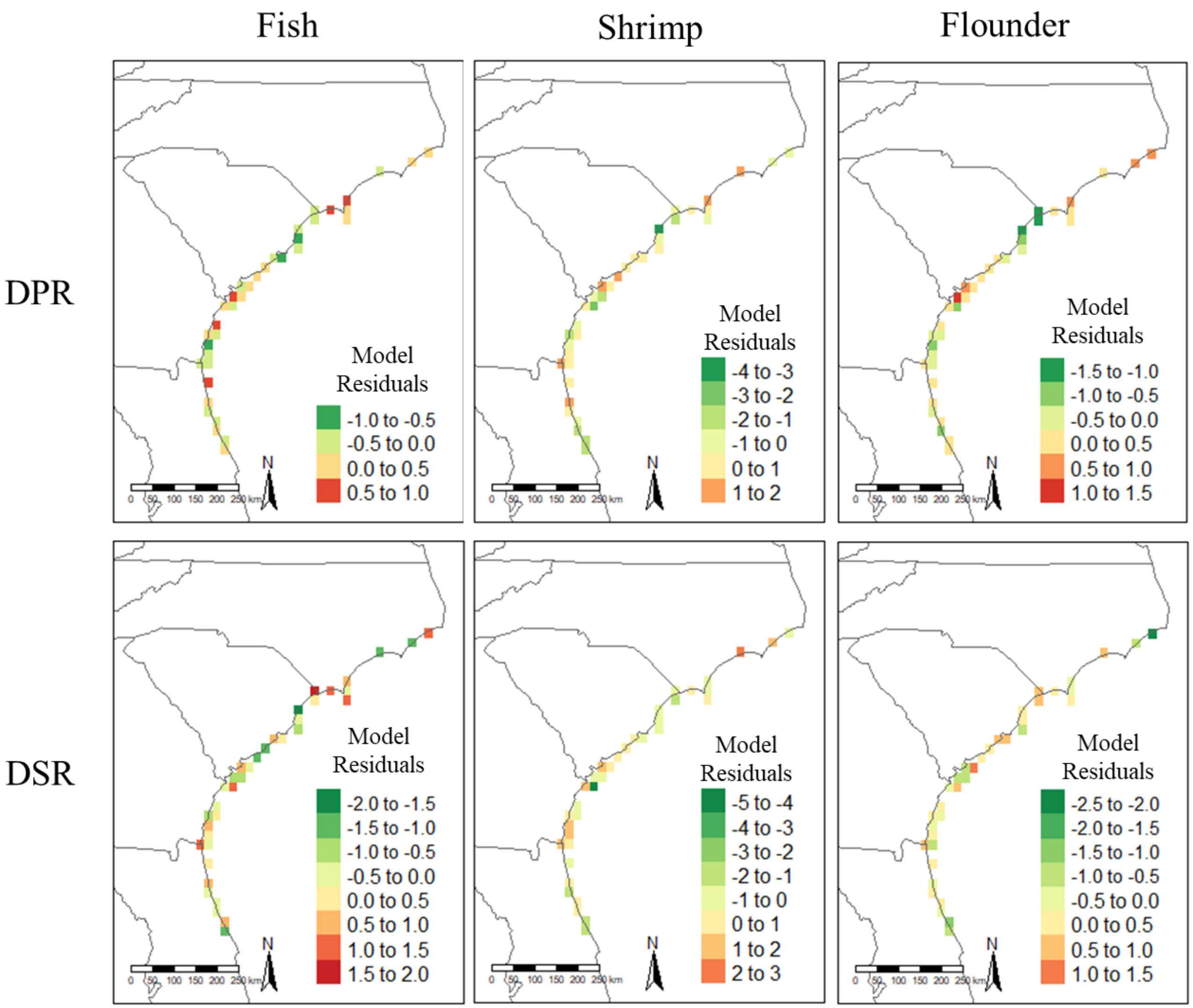


**Supplemental Figure S1**- Maps residuals of both the diversity-productivity (DPR) and diversity-stability (DSR) models for the three groups (all fish, shrimp, and flounder) throughout the 36 raster regions. Raster squares are approximately 22.2 km by 22.2km. See supplemental table 1 for model raw coefficients.
